# Protocols for fast simulations of protein structure flexibility using CABS-flex and SURPASS

**DOI:** 10.1101/694026

**Authors:** Aleksandra Badaczewska-Dawid, Andrzej Kolinski, Sebastian Kmiecik

## Abstract

Conformational flexibility of protein structures can play an important role in protein function. The flexibility is often studied using computational methods, since experimental characterization can be difficult. Depending on protein system size; computational tools may require large computational resources or significant simplifications in the modeled systems to speed-up calculations. In this work, we present the protocols for efficient simulations of flexibility of folded protein structures that use coarse-grained simulation tools of different resolutions: medium, represented by CABS-flex, and low, represented by SUPRASS. We test the protocols using a set of 140 globular proteins and compare the results with structure fluctuations observed in MD simulations, ENM modeling and NMR ensembles. As demonstrated, CABS-flex predictions show high correlation to experimental and MD simulation data, while SURPASS is less accurate but promising in terms of future developments.

## 1. Introduction

Molecular Dynamics (MD) and Elastic Network Models (ENM) are perhaps the most popular computational approaches in the studies of structural flexibility of biomolecules [1]. Both approaches are very effective in the study of local dynamics around well-defined, usually experimentally determined, structures that are used as the input. For MD studies, the size and complexity of the biomolecule may be a major limitation, while ENM approach may not give satisfactory results for structurally ambiguous regions [1]. The simulation alternative are coarse-grained (CG) protein models [2] which enable efficient modeling of much larger systems and/or longer processes than classical MD, and use more sophisticated interaction schemes than ENM techniques.

In this work, we present the protocols for prediction of protein structure fluctuations using CG protein models of medium-(CABS-flex) and low-resolution (SURPASS). Section 2 contains short description of CABS-flex [3,4] and SURPASS [5,6] simulation tools and their access links. Section 3 presents the protocols for fast simulations of protein structure flexibility. In order to test the protocols, we use the set of 140 globular proteins and compare the predictions of protein fluctuations with the data from all-atom MD, ENM (using DynOmics tool [7]) and NMR ensembles. The tests results are presented in section 4. In general, the CABS-flex method showed high correlation to experimental and MD simulation data. The low-resolution SURPASS was less accurate, particularly for proteins with low content of regular secondary structure or weak hydrophobic core, which is not surprising in the context of SURPASS design. Nevertheless, despite using very low-resolution of protein structure, SURPASS has showed accuracy on the level of the other methods for significant portion of the proteins from the test set. This is a very promising result in view of SURPASS potential use in studies of large protein systems. The advantageous feature of the tested protocols is their very low calculation cost, which is reduced to minutes in case of CABS-flex and seconds using SURPASS model (for proteins having several hundred residues and using a single CPU of average power).

## 2. Materials

### 2.1. CABS-flex method

CABS-flex method is dedicated for fast simulations of protein flexibility [3][4]. CABS-flex uses a well-established CABS coarse-grained model (reviewed elsewhere [2]) as a simulation engine and merges it with tools for all-atom and simulation analysis. The protein flexibility profiles produced by CABS Monte Carlo dynamics simulations were shown to be consistent with the protein flexibility of folded globular proteins seen in MD simulations [8,9], NMR ensembles [10] and also with various kinds of experimental data on protein folding mechanisms [11–14]. Moreover, CABS-flex method is successfully used in Aggrescan3D method [15–18] to predict the influence of protein flexibility on protein aggregation properties, and in CABS-dock method for simulations of protein flexibility during peptide molecular docking [19–22].

CABS-flex method is presently available as the CABS-flex 2.0 web server [3] (http://biocomp.chem.uw.edu.pl/CABSflex2) and the standalone package [4]. The package repository (available at https://bitbucket.org/lcbio/cabsflex/) contains online documentation, descriptions of the options, installation instructions and examples of usage. Note that CABS-flex standalone version runs on most Unix/Linux, Windows, MacOS systems and is also available as a Docker image. The following programs are also necessary:

1. *GFortran*, a freely redistributable Fortran compiler, required by CABS model (CABS-flex simulation engine)
2. *Python 2.7*, CABS-flex is a package using python 2.7 version
3. *Modeller* package [23], which is required by CABS-flex for all-atom reconstruction from C-alpha trace of CABS models

A thorough installation guide and CABS-flex issue tracker, which allows users to report any issues, can be found at CABS-flex repository.

### 2.2. SURPASS software

SURPASS [5,6] is a low-resolution coarse-grained model for efficient modeling of structure and dynamics of larger biomolecular systems. This model employs highly simplified representation of the protein structure and statistical potentials (see Figure 1). The concept of SURPASS representation is very simple and assumes averaging of short secondary structure fragments. The specific interaction model describes local structural regularities characteristic for most globular proteins. Despite its high simplification, SURPASS model reproduces reasonably well the basic structural properties of proteins and overcomes some limitations of coarse-grained moderate resolution models [5,6]. Reconstruction from SURPASS pseudo atoms to Cα-trace is possible using SUReLib algorithm (http://biocomp.chem.uw.edu.pl/tools/surpass).

**Figure 1.**
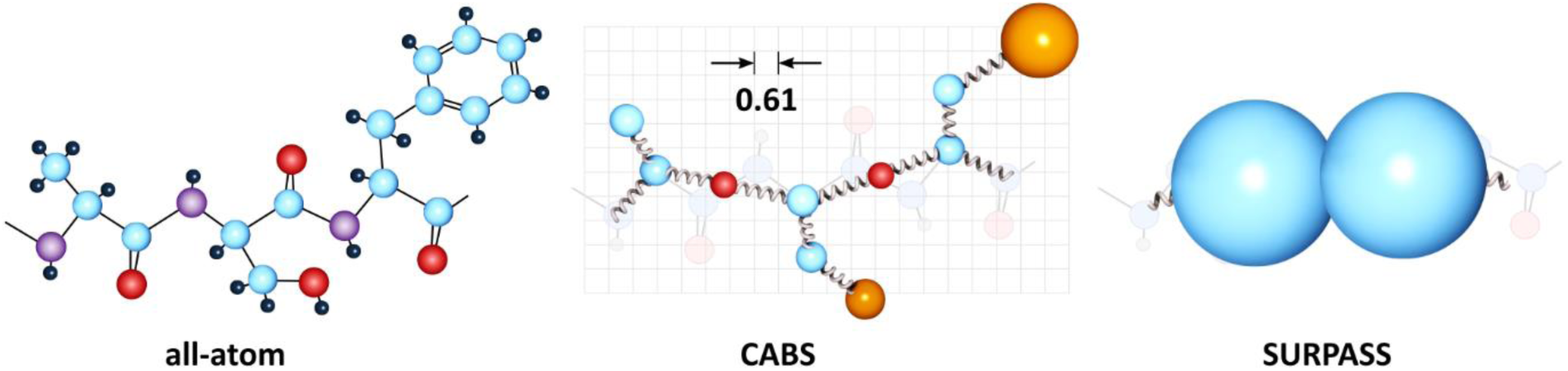
Example tripeptide in all-atom and coarse-grained (CG) representations. CABS CG model (used in CABS-flex package) and SURPASS CG model are presented. Single pseudo-atom of SURPASS replaces short (four residue long) secondary structure fragment. Despite the deep simplification of representation, the model reproduces basic structural properties of globular proteins.

The SURPASS software is available free of charge to academic use as a standalone program from the Laboratory website (http://biocomp.chem.uw.edu.pl/tools). Online SURPASS repository and documentation including installation instructions, options description and examples of use can also be found at https://bitbucket.org/lcbio/surpass/. Note that SURPASS is implemented in C++11 as a part of Bioshell 3.0 package and needs to be compiled (we recommend using the g++ ver. 4.9 compiler). After successful package compilation, you will find an executable program *surpass* in the bin directory. Calling a program with the *-h* option will display all currently available simulation settings. SURPASS software runs on most Unix/Linux, Windows, MacOS systems.

## 3. Methods

### 3.1. CABS-flex protocol for fast simulations of protein flexibility

The CABS-flex simulations of protein flexibility can be run using CABS-flex web server version [3] or the standalone package [4]. Here, we describe how to use the standalone package (a short information on using the web server is provided in Note 1). The only required CABS-flex input is a protein structure (in all-atom representation or C-alpha trace only). It may be provided as a file in PDB format, or just as PDB ID, which will be used by CABS-flex to download the appropriate file from the PDB database. To simulate only selected protein chains write appropriate chain symbols, e.g. „AC” after the colon sign. For example, to run CABS-flex for the protein with PDB ID 4w2o use one of the following commands:

~~~
*$ CABSflex -i 4w2o           #PDB ID variant for known protein structure
$ CABSflex -i PATH/structure_file.pdb #PATH is the localization of a structure_file.pdb on
your local system
$ CABSflex -i 4w2o:AC         #select chain ID if you don’t want to use all of them*
~~~

Running the CABS-flex protocol on the user’s local machine using the default simulation settings is as simple as in the example above. The default settings control the simulation parameters and distance restraints. The default CABS-flex settings were derived by Jamroz et al. [8] and provide a consensus picture of protein fluctuations with all-atom Molecular Dynamics in aqueous solution for globular proteins. Table 1 contains the exact set up of the default simulation settings. The detailed description of CABS-flex options is provided in CABS-flex repository at https://bitbucket.org/lcbio/cabsflex/. Below we comment only selected options. Modifications of these options may have some practical effects on the simulation outcome. The simulation length and number of models in the output trajectory are controlled by the values assigned to the set of *--mc* options (*-y, -s, -a*) (detailed description of the sampling procedure that is controlled by these options has been recently provided in the review by Ciemny et al. [24]). Using the *-t* option, user allows setting up the CABS simulation temperatures: at the beginning of the simulation (TINIT) and at the end of simulation (TFINAL). For example, the default setting “-t 1.4 1.4” introduce isothermal conditions at 1.4 temperature. This parameter may be used to increase or decrease the amplitude of protein fluctuations. The *--protein-restraints* option allow generating a set of binary distance restraints between Cα atoms. For example, the default setting of the *--protein-restraints* option: “ss2 3 3.8 8.0” make secondary structure elements more stable as compared to simulation without any restraints (see work by Jamroz et al. [8] for details). It is controlled by four parameters:

**Table 1.** Default CABS-flex simulation settings.

| option | short option | parameters | default value |
| --- | --- | --- | --- |
| --protein-restraints | -g | MODE GAP MIN MAX | ss2 3 3.8 8.0 |
| --temperature | -t | TINIT TFINAL | 1.4 1.4 |
| --replicas | -r | NUM | 1 |
| --mc-cycles | -y | NUM | 50 |
| --mc-steps | -s | NUM | 50 |
| --mc-annealing | -a | NUM | 20 |
| --protein-flexibility | -f | NUM or FILE | not applicable |

- MODE (default: ss2) which enables to select a subset of residues for which distance restraints will be generated, it can be between all residues [all], or only those belonging to secondary structure elements [ss2] or between residues from which at least one belong to secondary structure element [ss1];
- GAP (default: 3) specifies gap along the main chain (difference of indices) for two residues to be restrained;
- MIN (default: 3.8) and MAX (default: 8.0) define minimum and maximum restraint length in Angstroms between two residues to be restrained.

Additionally, protein flexibility (or rigidity, as defined in the CABS-flex web server [3]) can be modified with the *-f* option for selected protein residues (for the details see Note 2). By default, the simulation results are returned into 3 directories stored on the current/working path:

- *output_pdbs* – Cα-trace of initial structure (start.pdb), Cα-trace of trajectory (replica.pdb), 10 top models (all-atom) in separate PDB files and all models (Cα-trace) present in the 10 most dense clusters
- *output_data* – RMSD for each frame of trajectory comparing to reference structure (all_rmsds.txt); if no reference is given input structure is used as reference
- *plots* – data (.csv) and graphics (.svg) of Energy vs. RMSD and RMSF profile

### 3.1. SURPASS protocol for fast simulations of protein flexibility

The required SURPASS input is the protein structure of interest (all-atom or C-alpha trace only or SURPASS representation) provided in PDB file format and secondary structure assignment/prediction provided in ss2 file format. We recommend *dssp* as a method of assigning a secondary structure to a known protein structure or *psipred* as a method to predict a secondary structure (both approaches are presented in the Note 3). To run SURPASS using the protein with PDB ID 4w2o type the following command:

~~~
*$ ./surpass -in:pdb=4w2o.pdb -in:ss2=4w2o.ss2 -sample:t_start=0.2*
~~~

Running the SURPASS program on the user’s local machine is quite simple although it requires management from the command line. Table 1 contains the simulation settings with recommended values of parameters.

The length of the simulation and the number of models in the output trajectory is controlled by the values assigned to the set of *-sample:mc_* options. Using the *-sample:t_start* option user chooses the isothermal simulation scheme, which we recommend to study the local dynamics of folded protein structures. For other applications, that require enhanced sampling techniques, a simulated annealing scheme (additional options *-sample:t_end* and *-sample:t_steps*) or a Replica Exchange Monte Carlo sampling (use options *-sample:exchanges* and *-sample:replicas*) can be used. In contrast to CABS-flex, SURPASS does not automatically generate distance restraints based on the initial structure, but the user can load a two-column file with the indexes of the interacting residues. In this case, the information about the file with restraints should be provided in the configuration file (*surpass.wghts*, for the details see Note 4) of the SURPASS force field in the section concerning the *SurpassPromotedContact* energy component. The user can modify the strength of the given contacts in relation to new contacts created during the simulation. Changing the default settings allows the user to adjust the level of flexibility of the structure. Moreover, SURPASS provides many additional options that for example allow to load a reference structure for RMSD calculations, set the seed for random number generator or to modify the sampling scheme and many others (detailed descriptions are provided at https://bitbucket.org/lcbio/surpass/).

By default, the simulation results are returned into working directory. There are 4 categories of output files:

- *tra.pdb* – simulation trajectory in PDB file format (frames in SURPASS representation)
- *energy.dat* – scoring file with energy components
- *observers.dat* – for each frame contains *RgSquare, Elapsed_time* and *RMSD* to reference (or initial structure)
- *topology.dat* - for proteins containing beta-type secondary structure elements, assign a topological pattern for each frame

## 4. Case Studies

For the set of 140 globular proteins from the work of Jamroz et al. [10] we compared the predictions from CABS-flex, SURPASS, all-atom MD (deposited in the MoDEL database [25]), ENM (using DynOmics tool [7]) with NMR ensemble data. The input and reference structure for each protein was the first model from the NMR ensemble. CABS-flex, ENM and SURPASS simulations were run with default settings. For each protein from the test set, we obtained and analyzed the following data:

- 10 models from CABS-flex (by default, CABS-flex outputs 10 models obtained by structural clustering of 10’000 model’s trajectory)
- 40 models from DynOmics server [7] (two all-atom structures corresponding to each of 20 modes, for the details see Note 5)
- 100 models from SURPASS isothermal simulation (by default, all models from the trajectory)
- 10’000 models from MD simulation (data were taken from the MoDEL library [25] as Cα-only trajectory)
- at least 10 models (depending on protein) from NMR ensembles were taken from PDB database

For all simulation methods and NMR ensembles, the flexibility data were analyzed using root mean square fluctuation (RMSF) profiles and compared to the structural variability in NMR ensembles using Spearman’s rank correlation coefficient (rs_NMR_). In order to calculate these parameters, Theseus tool [26] was used to superimpose the models on the reference structure (the first model of the NMR ensemble, for the details see Note 6). Only Cα positions were used for structural alignment. In case of SURPASS, the pseudo atoms were superimposed on the reference structure converted into simplified representation. The residue fluctuation profiles were calculated according to the formula:

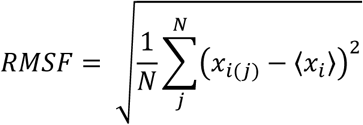

where *x*_*i*(*j*)_ denotes the position (coordinates) of the *i-*th Cα atom in the structure of the *j*-th model and ⟨*x*_*i*_⟩ denotes the averaged position of the *i*-th Cα atom in all models obtained by this method.

Spearman’s rank correlation coefficient between each method and NMR data was calculated as follows:

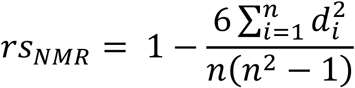

where *d*_*i*_ denotes the difference between the ranks of variables for the position of the *i*-th Cα atom, and *n* is the sum of Cα atoms in the system. Related ranks are taken into account as the arithmetic average of the ranks belonging to the same observations.

Table 1 shows RMSF and rs_NMR_ values (averaged over the entire set of 140 proteins). For each of the parameters in Table 1, the minimum (*min*), maximum (*max*) and average (*ave*) values are given. The lowest average RMSF fluctuations were observed for DynOmics server and the highest for SURPASS model (twice as high as for NMR). It should be noted that SURPASS simulations did not include any distance restraints so the structure was fully flexible. Moreover, the model currently does not have a component dependent directly on the amino acid sequence, which makes it less structurally rigid. Spearman’s rank correlation coefficient (*rs*_*NMR*_) is a measure of the statistical relationship between predicted residue fluctuation profiles and NMR ensemble data. For methods using medium or high resolution models, we observed a high (*rs*_*NMR*_>0.5) or very high (*rs*_*NMR*_>0.7, and in some cases > 0.9) correlation with NMR data (the highest *rs*_*NMR*_ of 0.96 was obtained using CABS-flex for 1P9C protein). In case of low-resolution SURPASS model, we noted the average level of correlation (*rs*_*NMR*_>0.3), although in some cases the level was high as for the other methods (e.g. for the proteins 1W9R or 2IQ3).

Table 2 contains a more detailed statistical analysis of Spearman’s rank correlation coefficient (*rs*_NMR_) dividing the results into two subsets of 1) average or weak correlation and 2) high and very high correlation. For each method, the percentage of deposits in a given range (counts) and the average value of the ratio in that range were given. Figure 2 shows correlation of RMSF profiles of different methods and NMR ensembles for 140 proteins from the test set. As demonstrated in Table 2 and Figure 2, most results significantly correlated with NMR ensembles were provided by DynOmics server (using ENM) and CABS-flex (more than 80% of proteins with an average correlation coefficient greater than 0.7).

**Table 1.**
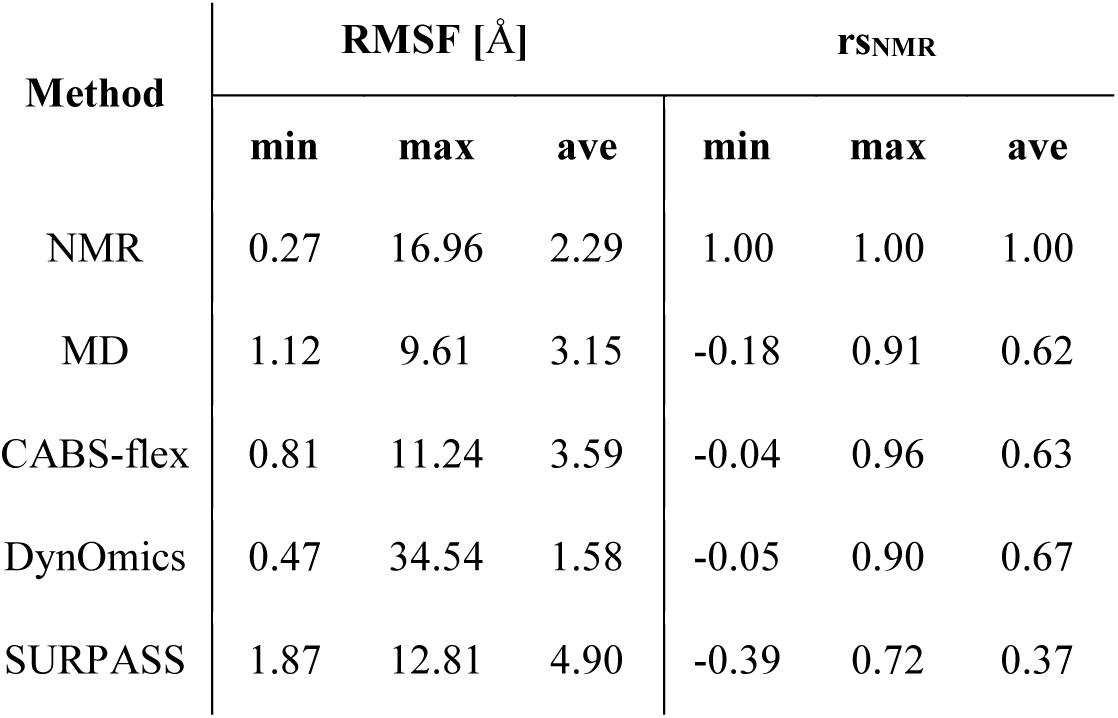
Average, minimum and maximum values of RMSF and rs_NMR_ for the benchmark set of 140 proteins We compare protein models obtained using MD, CABS-flex, DynOmics and SURPASS computational methods, and from NMR experimental data. RMSF is the averaged value of the fluctuation per residue. rs_NMR_ is the Spearman’s rank correlation coefficient calculated between the fluctuation profile for each method and the reference NMR data. Minimum (min), maximum (max) and mean (ave) values are given for both parameters.

| Method | RMSF [Å] | | | $r_{SNMR}$ | | |
| --- | --- | --- | --- | --- | --- | --- |
|  | min | max | ave | min | max | ave |
| NMR | 0.27 | 16.96 | 2.29 | 1.00 | 1.00 | 1.00 |
| MD | 1.12 | 9.61 | 3.15 | -0.18 | 0.91 | 0.62 |
| CABS-flex | 0.81 | 11.24 | 3.59 | -0.04 | 0.96 | 0.63 |
| DynOmics | 0.47 | 34.54 | 1.58 | -0.05 | 0.90 | 0.67 |
| SURPASS | 1.87 | 12.81 | 4.90 | -0.39 | 0.72 | 0.37 |

**Table 2.**
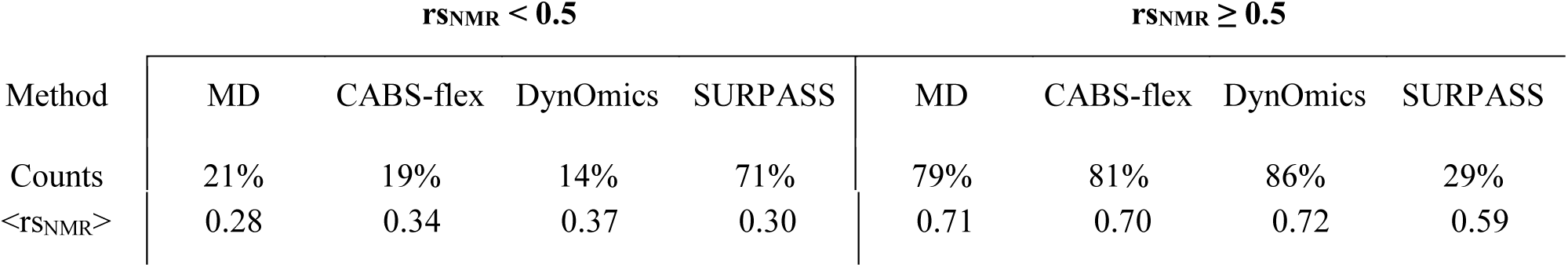
The mean of Spearman’s rank correlation coefficient (rs_NMR_) calculated between MD, CABS-flex, DynOmics, SURPASS methods and NMR data in two ranges of rs_NMR_ values: if less than 0.5 - average or weak correlation and if greater or equal to 0.5 - high or very high correlation.

| Method | $r_{SNMR} < 0.5$ | | | | $r_{SNMR} \geq 0.5$ | | | |
| --- | --- | --- | --- | --- | --- | --- | --- | --- |
|  | MD | CABS-flex | DynOmics | SURPASS | MD | CABS-flex | DynOmics | SURPASS |
| Counts | 21% | 19% | 14% | 71% | 79% | 81% | 86% | 29% |
| $\langle r_{\text{SNMR}} \rangle$ | 0.28 | 0.34 | 0.37 | 0.30 | 0.71 | 0.70 | 0.72 | 0.59 |

**Table 2.** Default SURPASS simulation settings.

| option | parameters | recommended value | description |
| --- | --- | --- | --- |
| -in:pdb | PATH/FILE | required, user provided | target structure |
| -in:ss2 | PATH/FILE | required, user provided | target secondary structure |
| -in_pdb:native | PATH/FILE | user provided | reference structure |
| -in:database | PATH | working dir | software database |
| -in:scfx | PATH/FILE | working dir | force field configuration |
| -sample:t_start | NUM (float) | 0.2 | isothermal temperature |
| -sample:mc_outer_cycles | NUM (int) | 100 | outer MC cycles |
| -sample:mc_inner_cycles | NUM (int) | 100 | inner MC cycles |

**Figure 2.**
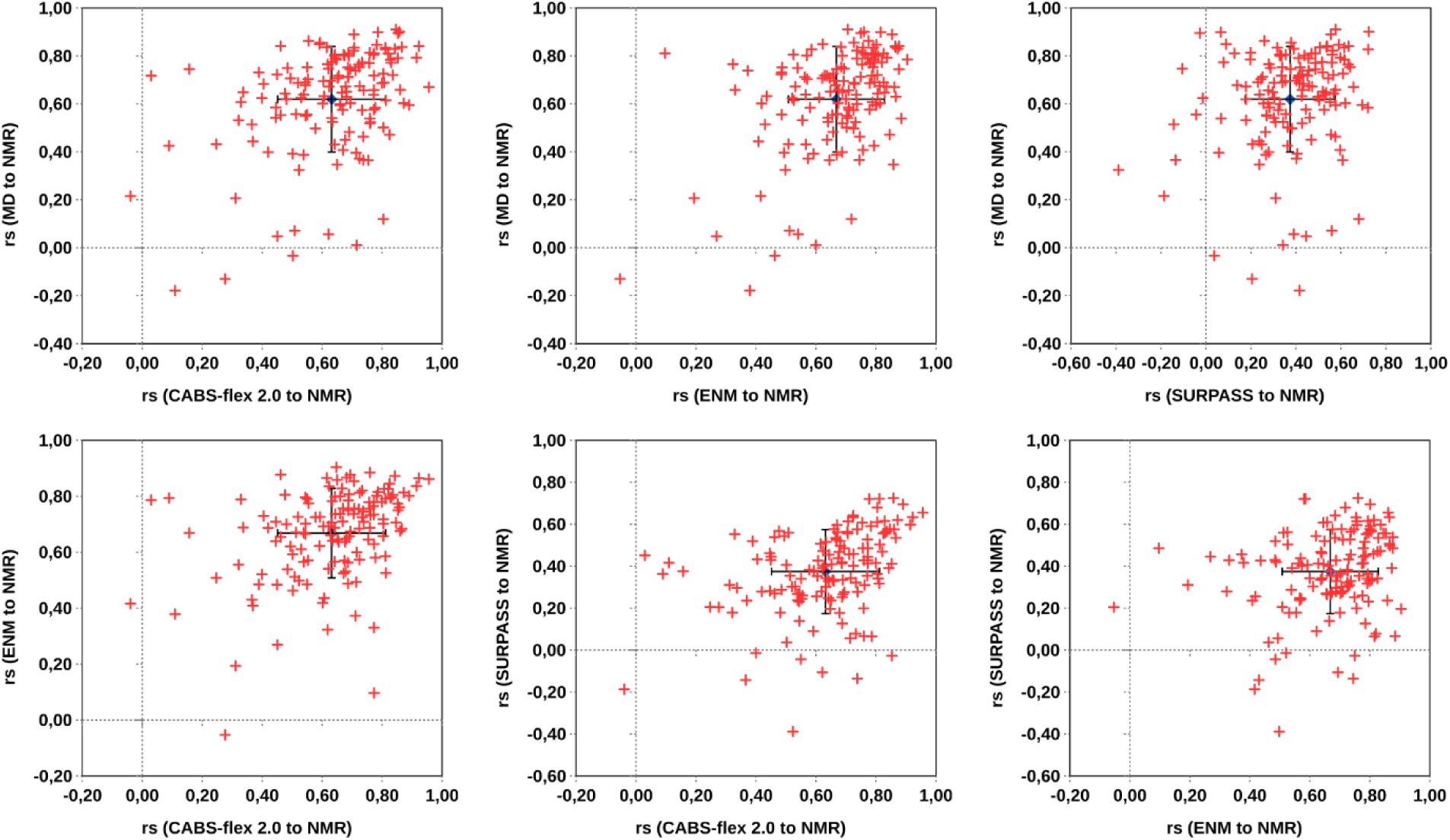
Spearman’s rank correlation coefficients of computational predictions using tested methods with NMR ensemble data (rs_NMR_) for 140 proteins. The graphs also show the mean value (blue dot) and standard deviation bars.

Figure 3 presents a comparison of the residue fluctuation profiles for selected proteins (profiles for entire benchmark set are shown in Note 7). 1IQ3 and 1W9R are an example of a protein for which all methods showed a high very high correlation to NMR data. 1SSG is an example of a small alpha protein for which all methods failed (see Note 7). The expected structural flexibility for this and several other cases (1DS9, 1EO0, 1K8B, 1PAW, 2RGF) is much higher than the reference NMR data, which may be caused by underestimation of structure fluctuations in NMR ensembles [27]. Interestingly, 1K5K, 1KGG, 1WAZ and 1RGF are also proteins for which the compliance of predicted fluctuation to experiment data was not high, but for all of them the best result was given by deeply simplified SURPASS model. Probably, the unrestrained mobility of SURPASS structures plays here a crucial role and this feature can be important for studies of highly flexible molecules.

**Figure 3.**
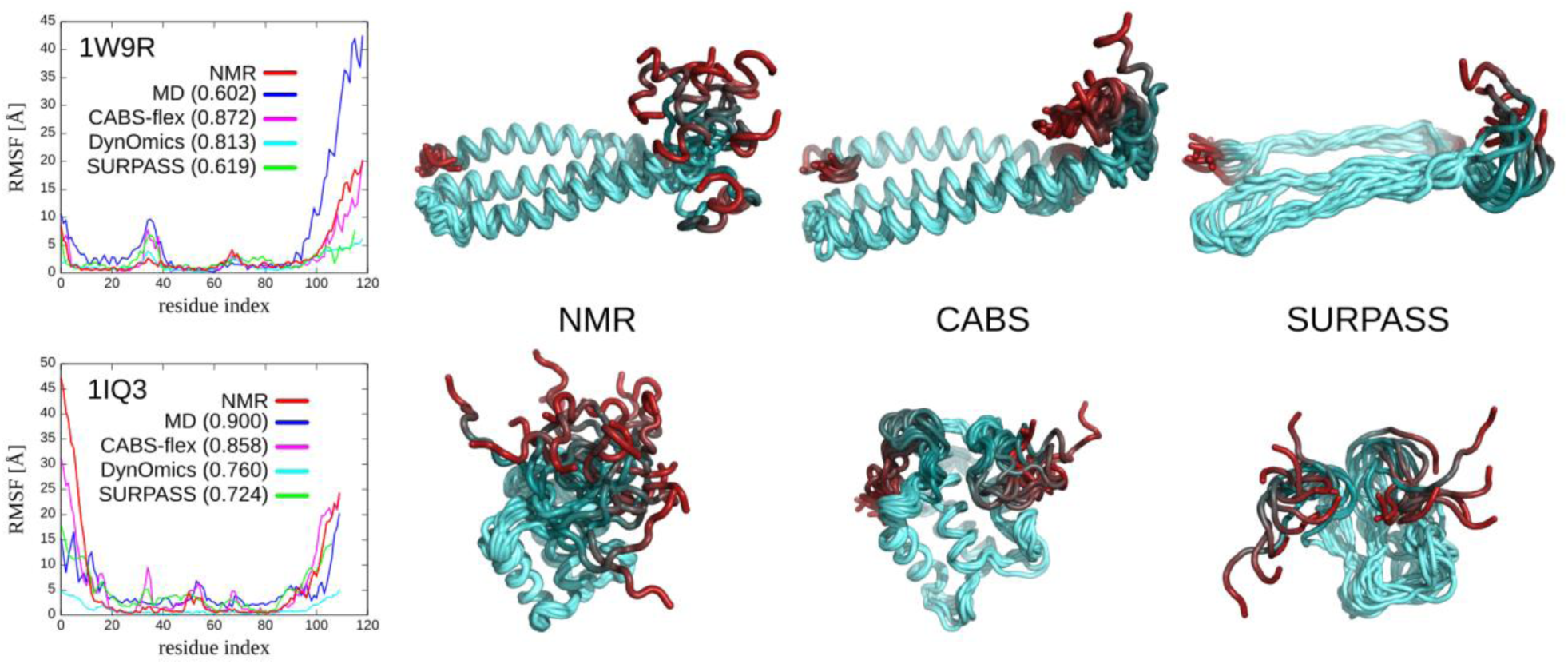
Residue fluctuation profiles for selected proteins (PDB code: 1W9R, 1IQ3) obtained using tested computational tools and from NMR ensembles. The curves on the plots were marked with colors: NMR - red, MD - blue, CABS-flex 2.0 - pink, DynOmics (ENM) - cyan, SURPASS - green. The Spearman’s rank correlation coefficient (rs_NMR_) is given in brackets. The right panel shows sets of protein models obtained by NMR, CABS and SURPASS methods.

For the studied set of 140 proteins the CABS-flex method showed high correlation to experimental data (Spearman’s rank correlation coefficient 0.7 for 81% of proteins). In most cases MD provided similar results, but for a few proteins MD predictions were much worse (1I35, 1OG7, 2HQI, 3CRD). In comparison to ENM-based tools (represented by the DynOmics tool in our tests), CABS-flex may be better suited for prediction of not obvious dynamic behavior, for example structure fluctuations within the well-defined secondary structural elements or other non-collective motions (see discussion in the review [1]). A low-resolution coarse-grained SURPASS model is less accurate than the other methods, which probably is related with a deep simplification of the interaction model, although the fluctuation profiles are quite realistic. This method showed a high or very high correlation to experimental NMR data for nearly 30% of proteins and an average for the rest. Much higher correlation to experimental fluctuation profiles was obtained for proteins with high contents of secondary structure elements (1W9R, 2IQ7). In a few cases the SURPASS model proved to be even better than the other methods (1KKG, 1WAZ). These results are very promising in terms of future SURPASS developments, that may focus on utilization of experimental data in the modeling process (for example in the form of distance restraints like in CABS-flex method). Finally, a particular advantage of the tested simulation protocols is their low computational cost (in the range of minutes for CABS-flex, or seconds for SURPASS using standard CPU). Both tested methods can be used as simulation engines of multiscale modeling protocols merging fast conformational sampling in coarse-grained resolution with more accurate modeling tools of higher resolution.

## 5. Notes

### Note 1

The CABS-flex web server [3] allows to make a similar computations as described in this work (obviously, the standalone package can be more useful for users interested in advanced options or in massive computations for many systems). Using the web server, user can provide an input structure by entering the ID of the protein from the Protein Data Bank (PDB) in the “PDB code” text box or by uploading a file in the pdb format from local hard drive in the “PDB file” box. The user can optionally provide protein chain(s) identifiers and project name, as well as an email address. Before the CABS-flex simulation begins, the server prepares a set of default distance restraints based on the input conformation. User can also create additional restraints or modify the initial ones by using tabs on the right marked as A and B in Figure 4. The third tab marked as C allows the user set few advanced simulation options. A preview of all additional options is given in the Figure 4 below. More details on using the server can be found in the web server publication or online server documentation at http://biocomp.chem.uw.edu.pl/CABSflex2/

**Figure 4.**
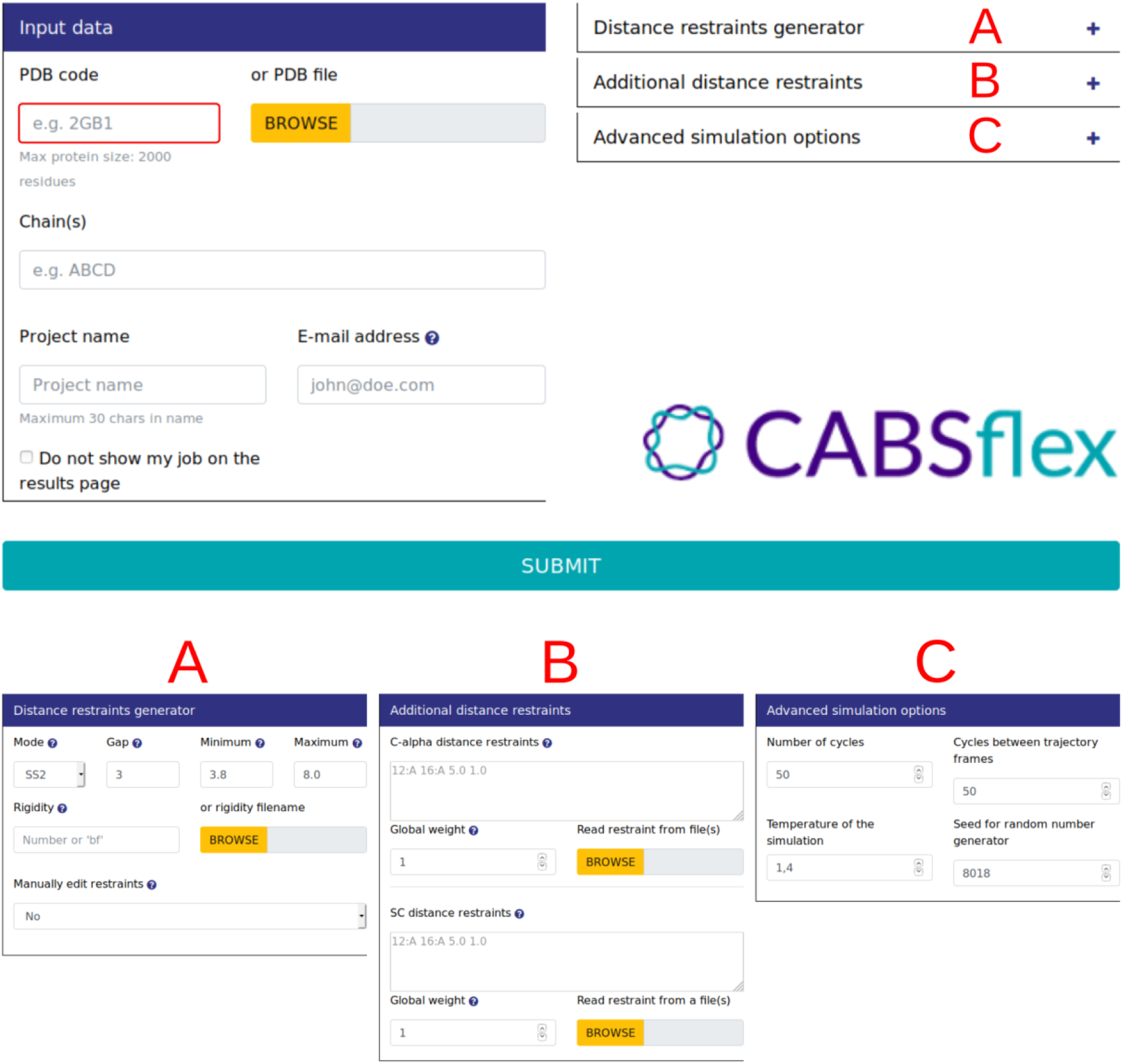
Interface of CABS-flex 2.0 web server [3]. The interface allows to modify or introduce simulation parameters and distance restraints.

**Figure 5.**
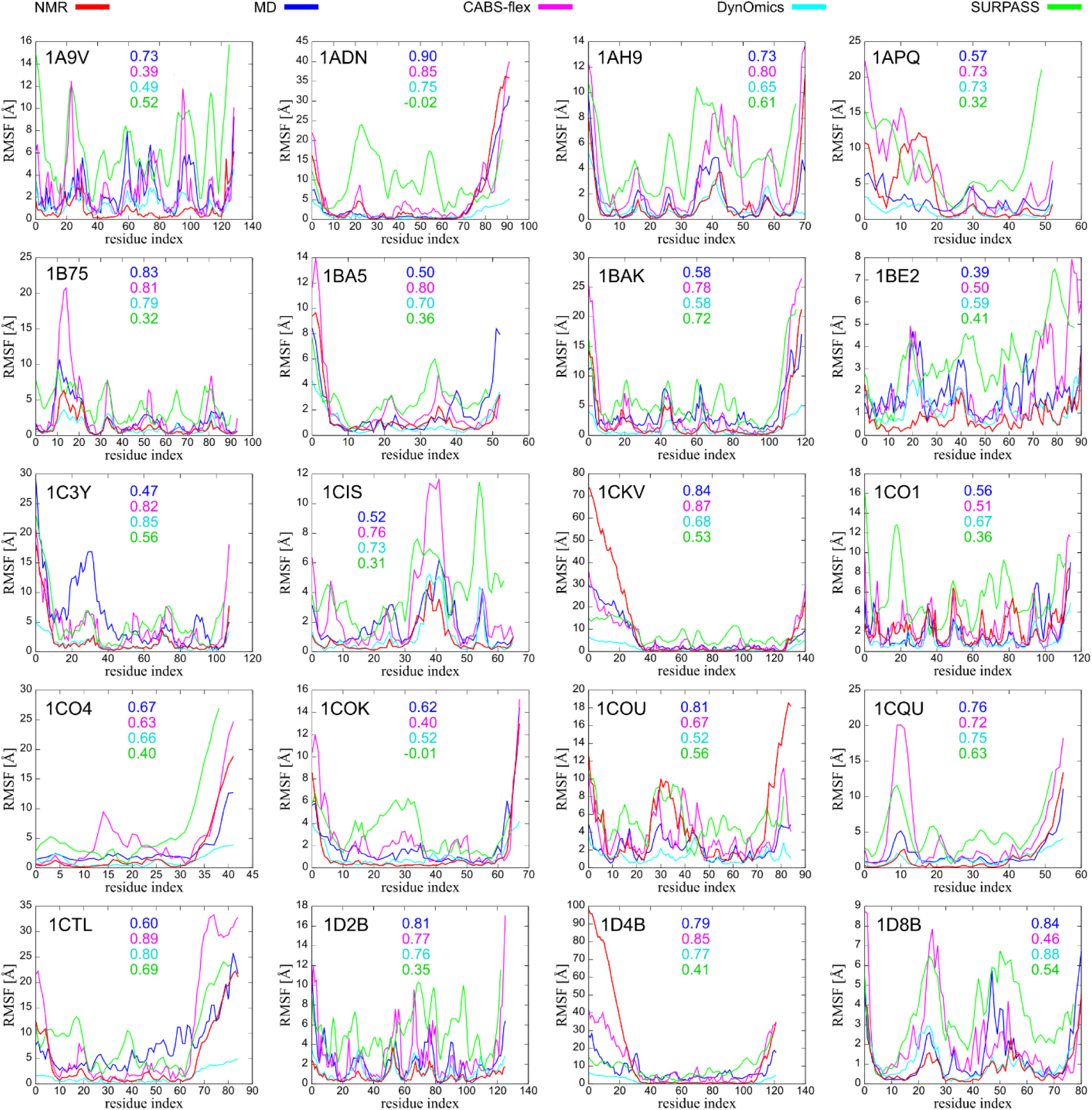
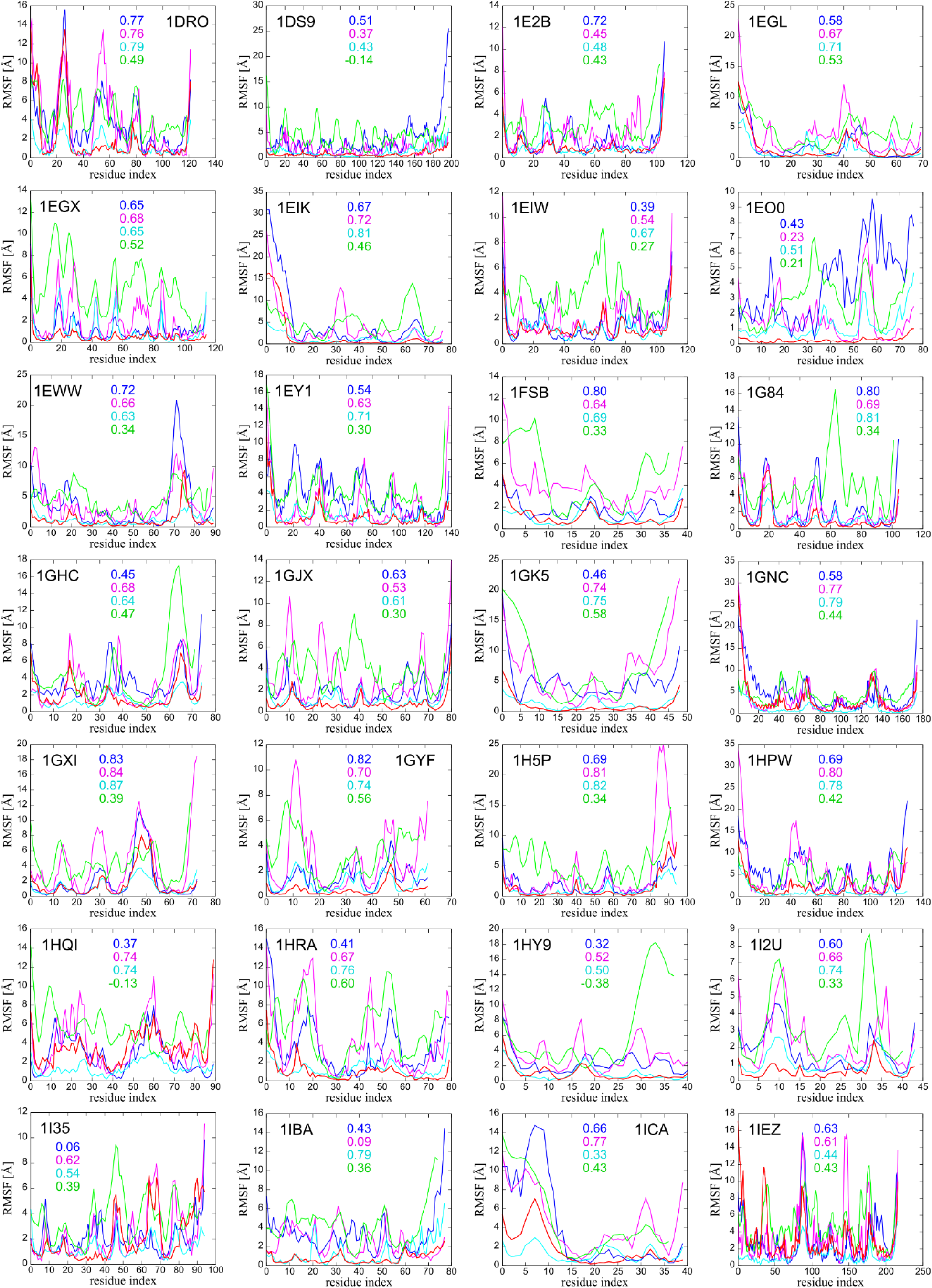
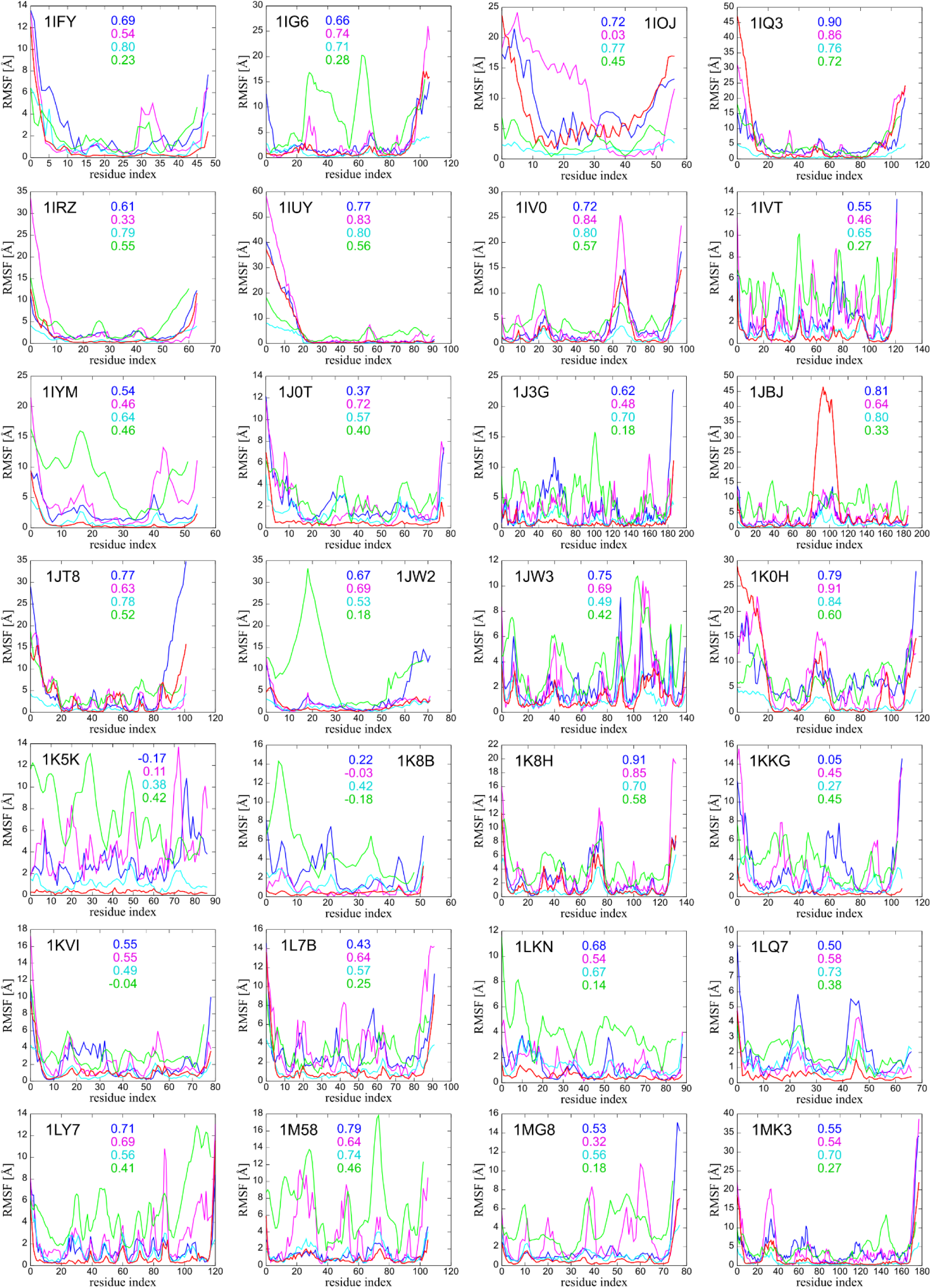
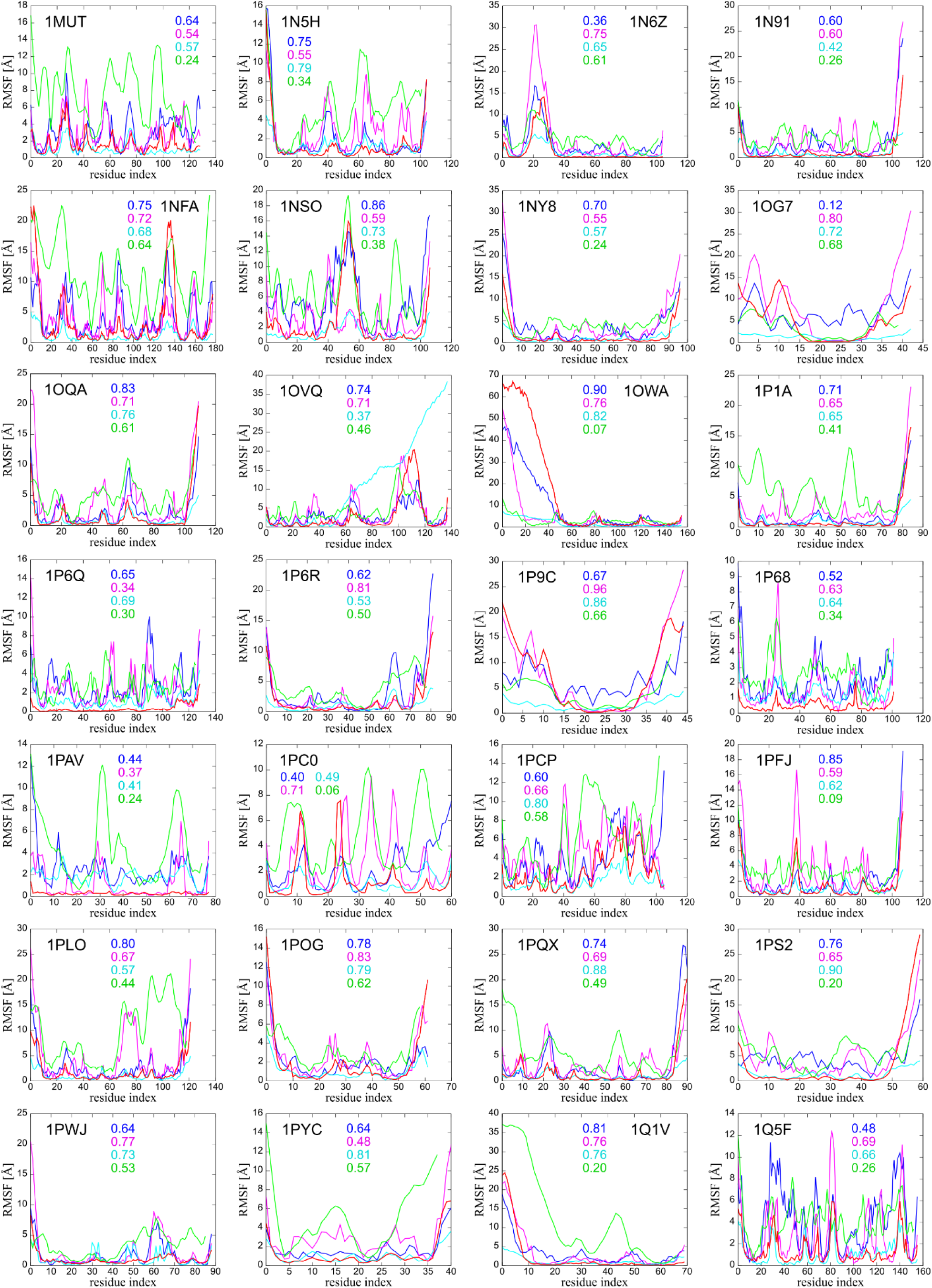
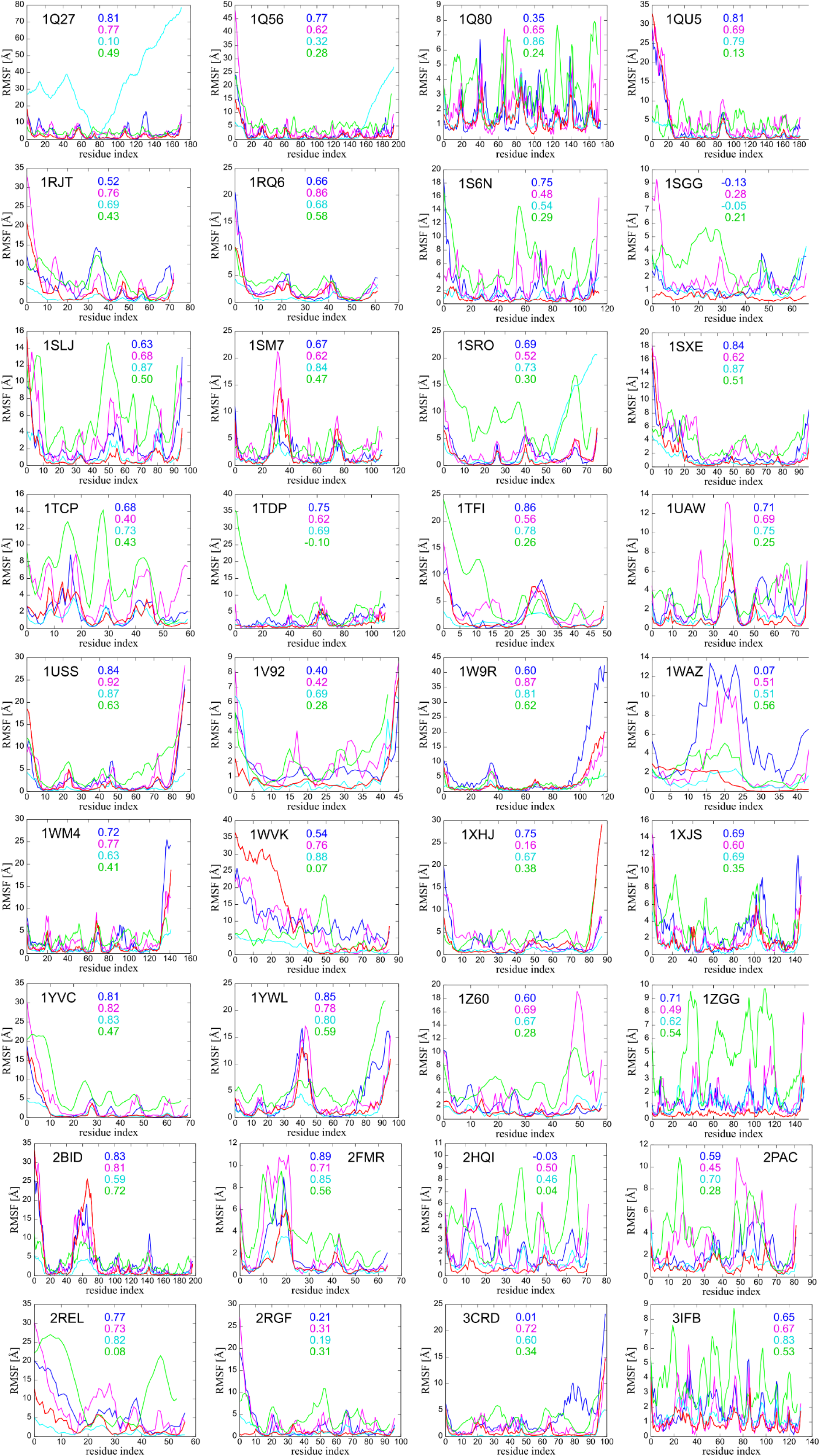
Residue fluctuation profiles for the benchmark set of 140 proteins obtained using tested computational tools (MD - blue, CABS-flex - pink, DynOmics - cyan, SURPASS - green) and from NMR ensembles (red). The numbers on the chart show the Spearman’s rank correlation coefficient between the computational method (see corresponding color) and the NMR data.

### Note 2

Except introducing or changing distance restraints, CABS-flex enables to modify structural flexibility of selected protein residues by using *-f* or *--protein-flexibility* option. To set up this particular simulation the user should prepare and load config-file telling CABS which fragment to modify and how much flexibility is needed. The configuration file, e.g. *4w2o.inp*, contains only one (or multiple) line:

~~~
*45:A - 51:A 0 #start – end of the fragment and flexibility value*
~~~

The flexibility value of selected protein residues can be modified as:

- 0 - fully flexible backbone,
- 1 - almost stiff backbone (default value, given appropriate number of protein restraints),
- >1 - increased stiffness,
- a positive real number - all protein residues will be assigned flexibility equal to this number,
- bf - flexibility for each residue is read from the beta factor column of the Cα atom in the PDB input file,
- bfi - each residue is assigned its flexibility based on the inverted beta factors stored in the input PDB file,
- <filename> - flexibility is read from file <filename> in the format of single residue entries, i.e. 12:A 0.75, or residue ranges, i.e. 12:A - 15:A 0.75

More details are provided in the CABS-flex repository available at https://bitbucket.org/lcbio/cabsflex/.

### Note 3

The idea of the SURPASS model is based on a specific averaging of the secondary structure fragments, therefore it is required to provide an assignment or prediction of the secondary structure in the ss2 format. If the spatial structure of the protein is known, we recommend using dssp to assign a secondary structure. The program is free available to download at https://swift.cmbi.umcn.nl/gv/dssp/DSSP_3.html. The program is executed from the command line with a simple command:

~~~
*$ mkdssp -i 4w2o.pdb -o 4w2o.dssp*
~~~

Then you can convert the output .*dssp* format file into required .*ss2* format using a ready-made program from the Bioshell package (*ap_dssp_to_ss2*), which you can find in the bin directory:

~~~
$ ./ *ap_dssp_to_ss2* 4w2o.dssp > 4w2o.ss2
~~~

If the protein structure is not known, we recommend using secondary structure predictor such as psipred, which is free available to download from github repository https://github.com/psipred/psipred. On the local machine, the program can be executed from the command line:

~~~
*$ ./runpsipred.local 4w2o.fasta* > *4w2o.horiz*
~~~

One of the default output files will be the predicted secondary structure in .*ss2* format. Note that only a protein sequence in .*fasta* format is required. Higher prediction accuracy can be achieved by using a consensus of various methods.

### Note 4

The knowledge-based SURPASS force field consists of several components that make up the total energy. Depending on the application, some potentials may be deactivated or the user may change their scaling factors, parameter values, or add distance restraints. The default configuration file, *surpass.wghts*, is located on the path ∼/data/forcefield/ in a local copy of the Bioshell package. The file can be copied to a working directory and modified as needed. Below you can find a preview of the config file with the description of the required parameters.

~~~
<u>./surpass.wghts</u>
*# R12 is a harmonic energy for pseudo-bonds*
**SurpassR12**   1.0 forcefield/local/R12_surpass.dat 0.001
*# R13 is a term that controls distance between i-th and (i*+*2) atoms*
**SurpassR13**    1.0 forcefield/local/R13_surpass.dat 0.001
*# R14 is a term that controls distance between i-th and (i*+*3) atoms*
**SurpassR14**    1.0 forcefield/local/R14_surpass.dat 0.001
*# R15 is a term that controls distance between i-th and (i*+*4) atoms*
**SurpassR15**    5.0 forcefield/local/R15_surpass.dat 0.001
*# A13 is a term that controls planar angle between i-th and (i*+*2) atoms*
**SurpassA13**    0.0 forcefield/local/A13_surpass.dat 0.001
*# SurpassHelixStifnessEnergy is a term that controls helix stifness*
**SurpassHelixStifnessEnergy** 5.0 -
*# SurpassCentrosymetricEnergy is forcing the presence of 50% of the residues at a predetermined distance*
**SurpassCentrosymetricEnergy** 1.0 -
*# SurpassLocalRepulsionEnergy is forcing the presence at most 6(E), 4(C), 2(H) residues in the local repulsion sphere*
**SurpassLocalRepulsionEnergy** 1.0 -
*# SurpassHydrogenBond calculate hydrogen bond energy only between atoms in B-sheets*
**SurpassHydrogenBond** 10.0 -
*# SurpassContactEnergy keeps excluded volume (repulsion) and contacts (attraction) parameters: weight, high_energy, low_energy, contact_shift*
**SurpassContactEnergy** 1.0 100.0 -0.5 0.01
*# SurpassPromotedContact promotes listed contacts
parameters: weight, high_energy, low_energy, contact_shift, restraints file, promote_weight*
**SurpassPromotedContact** 1.0 100.0 -0.5 0.01 PATH/surpass.contacts 5.0
~~~

### Note 5

DynOmics server (http://enm.pitt.edu/) enable to generate 2 all-atom structures along each of 20 modes at a given RMSD. Returned structures correspond to given (in Å) distance extremes of amplitude. For this purpose use the option: Molecular Motions → Full Atomic Structures for ANM-Driven Conformers → Motion along mode (1-20) with RMSD: (2Å) at Main result tab.

### Note 6

Theseus (https://theobald.brandeis.edu/theseus/) is an efficient program for superpositioning multiple macromolecular structures using the method of maximum likelihood. The program is executed from the command line with a simple command:

~~~
*$ ./theseus reference.pdb target.pdb*
~~~

Among the output files there are two in PDB format:

- theseus_ave.pdb - artificially averaged structure (single structure)
- theseus_sup.pdb - original structures imposed on the reference (multiple structures)

Note that in our studies, the reference structure for both the imposition and all calculations was the first model from NMR ensemble, not the “artificially” averaged structure generated by Theseus.

### Note 7

## Acknowledgments

A.B-D, A.K., and S.K. received funding from NCN Poland, Grant MAESTRO2014/14/A/ST6/00088.

## References

1. Kmiecik, S.; Kouza, M.; Badaczewska-Dawid, A. E.; Kloczkowski, A.; Kolinski, A. Modeling of Protein Structural Flexibility and Large-Scale Dynamics: Coarse-Grained Simulations and Elastic Network Models. Int. J. Mol. Sci. 2018, 19, E3496, doi:10.3390/ijms19113496.

2. Kmiecik, S.; Gront, D.; Kolinski, M.; Wieteska, L.; Dawid, A. E.; Kolinski, A. Coarse-Grained Protein Models and Their Applications. Chem. Rev. 2016, 116, doi:10.1021/acs.chemrev.6b00163.

3. Kuriata, A.; Gierut, A. M.; Oleniecki, T.; Ciemny, M. P.; Kolinski, A.; Kurcinski, M.; Kmiecik, S. CABS-flex 2.0: A web server for fast simulations of flexibility of protein structures. Nucleic Acids Res. 2018, doi:10.1093/nar/gky356.

4. Kurcinski, M.; Oleniecki, T.; Ciemny, M. P.; Kuriata, A.; Kolinski, A.; Kmiecik, S. CABS-flex standalone: a simulation environment for fast modeling of protein flexibility. Bioinformatics 2018, 35, 694–695, doi:10.1093/bioinformatics/bty685.

5. Dawid, A. E.; Gront, D.; Kolinski, A. SURPASS Low-Resolution Coarse-Grained Protein Modeling. J. Chem. Theory Comput. 2017, 13, doi:10.1021/acs.jctc.7b00642.

6. Dawid, A. E.; Gront, D.; Kolinski, A. Coarse-Grained Modeling of the Interplay between Secondary Structure Propensities and Protein Fold Assembly. J. Chem. Theory Comput. 2018, 14, 2277–2287, doi:10.1021/acs.jctc.7b01242.

7. Li, H.; Chang, Y. Y.; Lee, J. Y.; Bahar, I.; Yang, L. W. DynOmics: Dynamics of structural proteome and beyond. Nucleic Acids Res. 2017, 45, W374–380, doi:10.1093/nar/gkx385.

8. Jamroz, M.; Orozco, M.; Kolinski, A.; Kmiecik, S. Consistent view of protein fluctuations from all-atom molecular dynamics and coarse-grained dynamics with knowledge-based force-field. J. Chem. Theory Comput. 2013, doi:10.1021/ct300854w.

9. Jamroz, M.; Kolinski, A.; Kmiecik, S. CABS-flex: Server for fast simulation of protein structure fluctuations. Nucleic Acids Res. 2013, doi:10.1093/nar/gkt332.

10. Jamroz, M.; Kolinski, A.; Kmiecik, S. CABS-flex predictions of protein flexibility compared with NMR ensembles. Bioinformatics 2014, doi:10.1093/bioinformatics/btu184.

11. Kurcinski, M.; Kolinski, A.; Kmiecik, S. Mechanism of folding and binding of an intrinsically disordered protein as revealed by ab initio simulations. J. Chem. Theory Comput. 2014, doi:10.1021/ct500287c.

12. Kmiecik, S.; Kolinski, A. Characterization of protein-folding pathways by reduced-space modeling. Proc. Natl. Acad. Sci. 2007, 104, 12330–12335, doi:10.1073/pnas.0702265104.

13. Kmiecik, S.; Kolinski, A. Folding pathway of the B1 domain of protein G explored by multiscale modeling. Biophys. J. 2008, doi:10.1529/biophysj.107.116095.

14. Kmiecik, S.; Gront, D.; Kouza, M.; Kolinski, A. From coarse-grained to atomic-level characterization of protein dynamics: Transition state for the folding of B domain of protein A. J. Phys. Chem. B 2012, 116, doi:10.1021/jp301720w.

15. Zambrano, R.; Jamroz, M.; Szczasiuk, A.; Pujols, J.; Kmiecik, S.; Ventura, S. AGGRESCAN3D (A3D): Server for prediction of aggregation properties of protein structures. Nucleic Acids Res. 2015, doi:10.1093/nar/gkv359.

16. Kuriata, A.; Iglesias, V.; Pujols, J.; Kurcinski, M.; Kmiecik, S.; Ventura, S. Aggrescan3D (A3D) 2.0: prediction and engineering of protein solubility. Nucleic Acids Res. 2019, gkz321, doi:10.1093/nar/gkz321.

17. Gil-Garcia, M.; Bañó-Polo, M.; Varejão, N.; Jamroz, M.; Kuriata, A.; Díaz-Caballero, M.; Lascorz, J.; Morel, B.; Navarro, S.; Reverter, D.; Kmiecik, S.; Ventura, S. Combining Structural Aggregation Propensity and Stability Predictions To Redesign Protein Solubility. Mol. Pharm. 2018, 15, 3846–3859, doi:10.1021/acs.molpharmaceut.8b00341.

18. Kuriata, A.; Iglesias, V.; Kurcinski, M.; Ventura, S.; Kmiecik, S. Aggrescan3D standalone package for structure-based prediction of protein aggregation properties. Bioinformatics 2019, doi:10.1093/bioinformatics/btz143.

19. Kurcinski, M.; Jamroz, M.; Blaszczyk, M.; Kolinski, A.; Kmiecik, S. CABS-dock web server for the flexible docking of peptides to proteins without prior knowledge of the binding site. Nucleic Acids Res. 2015, doi:10.1093/nar/gkv456.

20. Blaszczyk, M.; Kurcinski, M.; Kouza, M.; Wieteska, L.; Debinski, A.; Kolinski, A.; Kmiecik, S. Modeling of protein-peptide interactions using the CABS-dock web server for binding site search and flexible docking. Methods 2016, doi:10.1016/j.ymeth.2015.07.004.

21. Ciemny, M. P.; Kurcinski, M.; Kozak, K.; Kolinski, A.; Kmiecik, S. Highly flexible protein-peptide docking using cabs-dock. Methods Mol. Biol. 2017, 1561, 69–94, doi:10.1007/978-1-4939-6798-8_6.

22. Kurcinski, M.; Ciemny, M. P.; Oleniecki, T.; Kuriata, A.; Badaczewska-Dawid, A. E.; Kolinski, A.; Kmiecik, S. CABS-dock standalone: a toolbox for flexible protein-peptide docking. Bioinformatics 2019, doi:10.1093/bioinformatics/btz185.

23. Webb, B.; Sali, A. Comparative protein structure modeling using MODELLER. Curr. Protoc. Bioinforma. 2016, 54, 5.6.1-5.6.37, doi:10.1002/cpbi.3.

24. Ciemny, M. P.; Badaczewska-Dawid, A. E.; Pikuzinska, M.; Kolinski, A.; Kmiecik, S. Modeling of Disordered Protein Structures Using Monte Carlo Simulations and Knowledge-Based Statistical Force Fields. Int. J. Mol. Sci. 2019, 20, E606, doi:10.3390/ijms20030606.

25. Meyer, T.; D’Abramo, M.; Hospital, A.; Rueda, M.; Ferrer-Costa, C.; Pérez, A.; Carrillo, O.; Camps, J.; Fenollosa, C.; Repchevsky, D.; Gelpí, J. L.; Orozco, M. MoDEL (Molecular Dynamics Extended Library): A Database of Atomistic Molecular Dynamics Trajectories. Structure 2010, 18, 1399–1409, doi:10.1016/j.str.2010.07.013.

26. Theobald, D. L.; Wuttke, D. S. THESEUS: Maximum likelihood superpositioning and analysis of macromolecular structures. Bioinformatics 2006, 22, 2171–2172, doi:10.1093/bioinformatics/btl332.

27. Spronk, C. A. E. M.; Nabuurs, S. B.; Bonvin, A. M. J. J.; Krieger, E.; Vuister, G. W.; Vriend, G. The precision of NMR structure ensembles revisited. J. Biomol. NMR 2003, 25, 225–34, doi:10.1023/A:1022819716110.

